# Shapes and Dimensions of Blood Clots Affect the Rate and Extent of Platelet-Driven Clot Contraction and Will Determine the Outcomes of Thrombosis

**DOI:** 10.1101/2025.03.30.646236

**Authors:** Rafael R. Khismatullin, Alina I. Khabirova, Shakhnoza M. Saliakhutdinova, Rustem I. Litvinov, John W. Weisel

**Author notes:** Correspondence: Prof. John W. Weisel, Department of Cell and Developmental Biology University of Pennsylvania School of Medicine 421 Curie Blvd., BRB II/III, Room 1154 Philadelphia, PA 19104-6058, USA.

## Abstract

**Background:** Contraction (retraction, shrinkage) of hemostatic blood clots and obstructive thrombi is an important pathophysiological process. To mimic thrombi of various locations, we sought to determine whether the initial shape and dimensions of blood clots affect the rate and extent of clot contraction.

**Methods:** Thrombin-induced clots were formed in 0.3-5.7 ml samples of citrated human blood or platelet-rich plasma in a cylinder, cuboid, or flat chamber. The clots were allowed to contract at 37°C for 60 minutes with shrinkage tracked photographically. Following complete contraction, the physiologically most relevant cylindrical clots of various initial volumes (0.5 ml or 1.5 ml) were analyzed with scanning electron microscopy for composition and spatial non-uniformity with the emphasis on compressed polyhedral erythrocytes (polyhedrocytes).

**Results:** With the same volumes studied, the rates and final extents of contraction of whole blood clots were shape-dependent in the following order: flat > cuboid = cylindrical. Irrespective of the shape, the initially smaller clots always contracted to a larger extent. Unlike clots in whole blood, the platelet-rich plasma clots contracted almost independently of the clot volumes and shapes studied, indicating a key role of erythrocytes. The smaller blood clots with a higher extent of contraction contained more relative volume fraction of erythrocytes, especially compressed polyhedrocytes, due to tight packing and a decrease in the intercellular space. Unlike the smaller clots, the larger clots were not clearly segregated into distinct layers, reflecting incomplete spatial redistribution of blood clot components typical for contraction.

**Conclusions:** Contraction of blood clots depends on their shape and size. The smaller and larger clots have distinct size-dependent rates and extents of contraction as well as degrees of structural non-uniformity, reflecting different spatial gradients of compressive stresses. The physiological relevance of these findings is related to the variable geometry and size of intravascular blood clots and thrombi.

## Introduction

Blood clot contraction or retraction is the active shrinkage of a clot occurring both *in vitro* and *in vivo* that makes the clot more compact, stiff, and less permeable [*1; 2*]. Contraction of a hemostatic blood clot helps to seal the site of injury, preventing blood loss [*3; 4*] and improving wound healing [5], while contraction of an obstructive thrombus within a vessel restores local blood flow past the otherwise obstructive thrombus [*6; 7*]. In addition, clot contraction *in vivo* reduces the risk of thrombotic embolization [*8; 9*] and alters clot susceptibility to fibrinolysis [10].

Activated platelets are the driving force of blood clot contraction; their intracellular adenosine triphosphate (ATP)-dependent actomyosin machinery generates traction forces that are transmitted outside to fibrin fibers via integrin αIIbβ3, causing compaction of the fibrin network and entire clot [11–13]. In the course of contraction, blood clots undergo profound structural rearrangements and acquire at least two major morphological features: i) spatial redistribution of clot components, such that red blood cells (RBCs) are accumulated and packed tightly in the interior, while a meshwork of fibrin and platelets accumulates at the clot periphery, and ii) gradual deformation of RBCs from their customary biconcave to a compressed polyhedral shape [*1; 14*]. These morphological signatures of clot contraction revealed initially *in vitro*, were also found in *ex vivo* thrombi of various origins, indicating that contraction of blood clots and thrombi is a pathophysiological process that has important clinical implications [*7-9; 15-18*].

Because blood clot contraction is a multifactorial process, the rate and extent of blood clot shrinkage can be modulated by a number of factors [19], such as platelet counts and platelet functionality [19–23]; the counts, mechanical properties, and functional states of RBCs [19] and leukocytes [24]; and the mass and mechanical properties of fibrin, including factor XIIIa-catalyzed fibrin cross-linking [*19*; *25; 26*]. Clot contraction is partially impaired in (pro)thrombotic conditions of various etiologies [*7; 8; 27-32*], which is due to hypercoagulability and systemic thrombinemia, leading to continuous partial platelet activation in the circulation followed by their exhaustion and dysfunction, including reduced contractility [*33*; *34*].

Despite many studies on the pathophysiological relevance of blood clot contraction and its modulation, it remains unknown whether this biomechanical process depends on the initial clot geometry and size. This aspect seems to be of great clinical importance as *in vivo* thrombi have variable geometry and size determined by the geometrically diverse blood vessels or extravascular space in bleeding. From live imaging, surgical findings, and autopsy, thrombi vary in shape in size from flattened parietal short clots in some arteries to long occlusive cylindrical clots in veins of lower limbs [35–37]. Many studies demonstrated that the thrombus location and conditions of its formation, as well as vessel diameter, play a crucial role in the course and outcomes of thrombosis. In particular, arterial thrombosis is usually associated with vascular damage, high shear stress with predominant platelet activation, and platelet-vessel wall interaction [*38; 39*], whereas venous thrombosis most often results from a combination of hypercogulability, blood stasis and vascular damage known as Virchov’s triad [*40; 41*]. Thus, different pathophysiological mechanisms of formation of thrombi at various locations (large, medium or small arteries or veins) cause diversity of their macro- and microscopic morphology that is tightly associated with biological and physical properties, such as obstructiveness, lytic and mechanical stability, permeability, etc. The goal of this work was to determine the relationships between the volume and shape of blood clots and their ability to undergo biomechanical and structural remodeling known as clot contraction.

## Methods

Details on the collection and fractionation of human blood samples; formation of *in vitro* blood and platelet-rich plasma (PRP) clots of various shapes and volumes; types of plastic or glass chambers used to form clots and follow their shrinkage; time-lapse imaging and measuring clot volume during contraction; scanning electron microscopy (SEM) of blood clots and quantitative SEM image analysis; as well as statistical analyses are provided in the Data Supplement. The authors declare that all supporting data are available within the article and its Data Supplement.

Briefly, thrombin-induced clots from human citrated whole blood or PRP were formed in 0.3-5.7 ml samples in a cylinder, cuboid, or flat chamber, and were allowed to shrink at 37°C for 60 minutes (Fig. 1). To quantify the kinetics of clot contraction, time-dependent changes in the volume of contracting blood or plasma clots were determined from photographic images taken every 5 minutes during 1 hour following clot formation. The physiologically most relevant cylindrical fully contracted clots of various initial volumes were analyzed using SEM for composition and spatial non-uniformity (Fig. 2), with the emphasis on redistribution of fibrin and compressed red blood cells as the morphological features and quantitative measures of the extent of clot contraction (Fig. 3).

**Figure 1.**
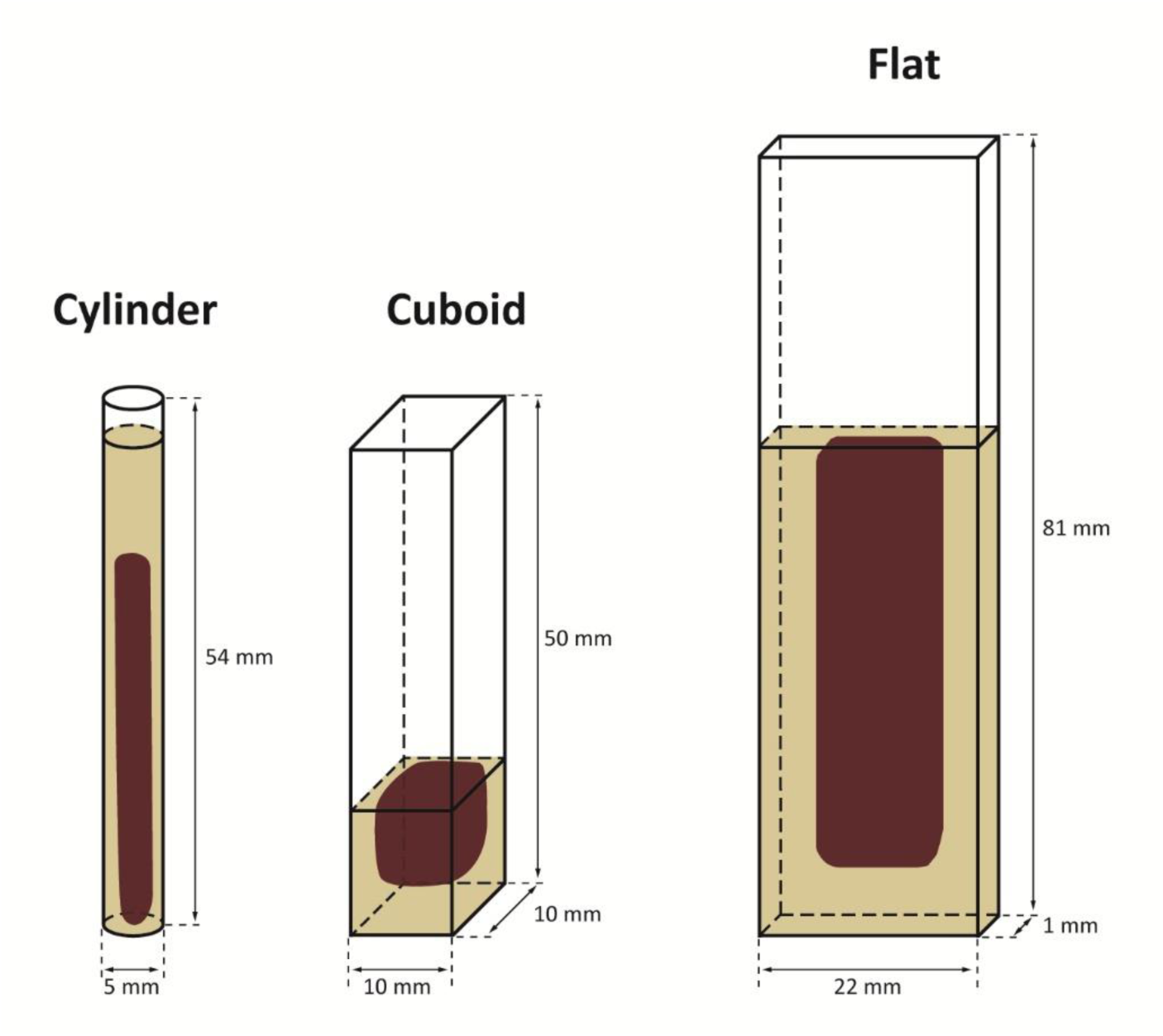
Schematic diagrams (not to scale) showing examples of the types of chambers used to form whole blood or PRP clots of various shapes and size (exemplified with 1.5-ml whole blood clots after contraction).

**Figure 2.**
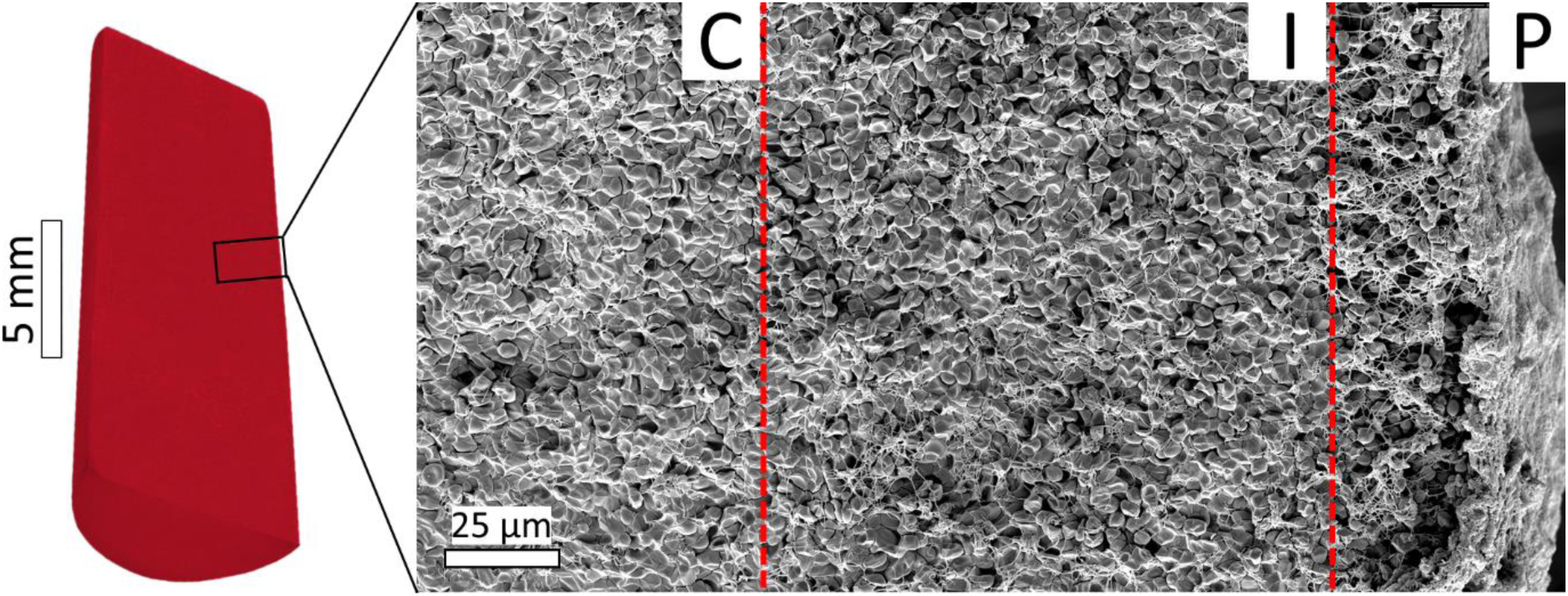
A schematic cylindrical blood clot sectioned longitudinally through the middle plane (*left*) and panoramic (technology of our scanning electron microscope to stitch together hundreds of adjacent images) scanning electron micrograph (*right*) from an inner portion of contracted 0.5-ml cylindrical blood clot with the central (*C*), intermediate (*I*), and peripheral (*P*) layers characterized by distinct packing densities.

**Figure 3.**
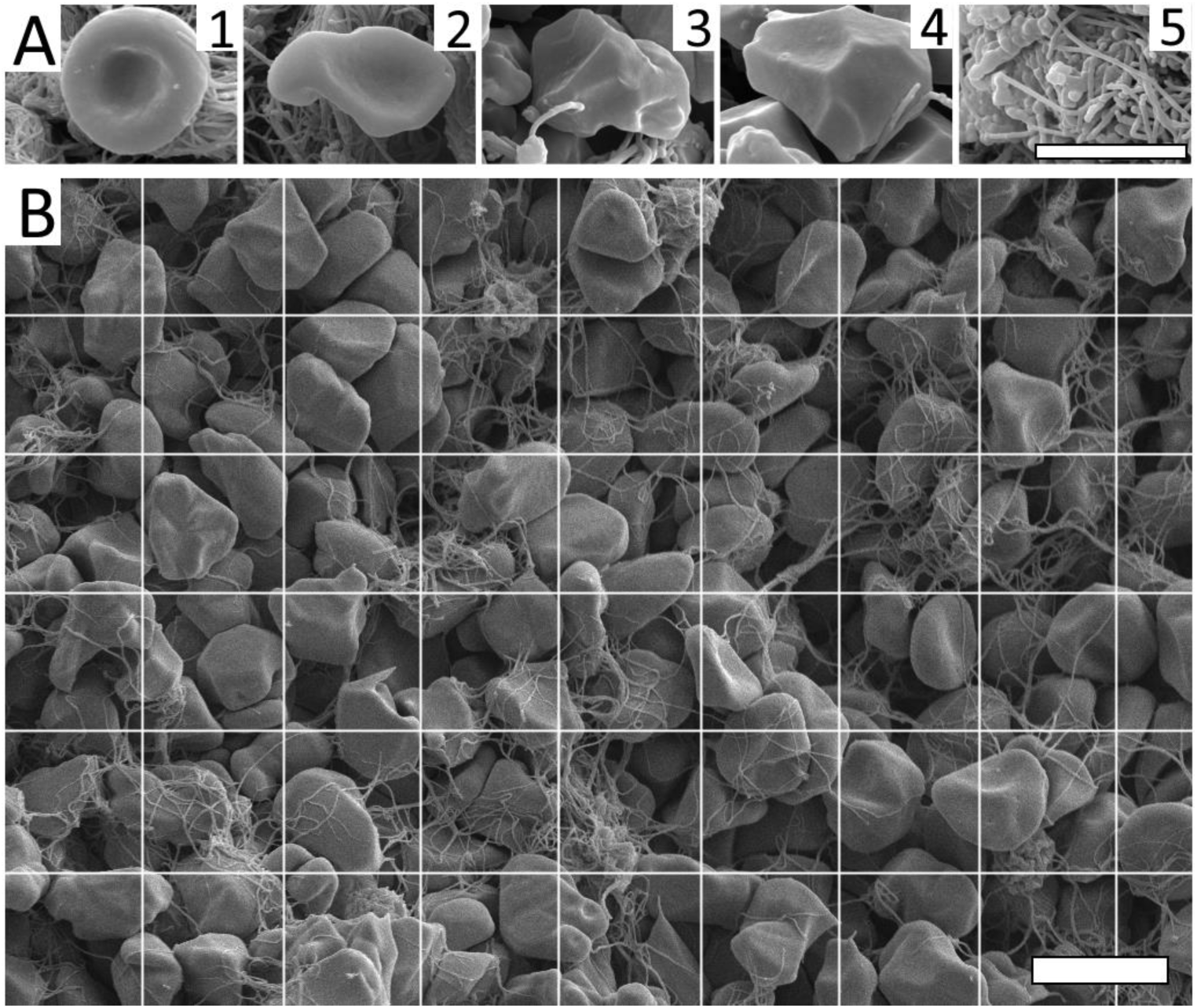
Imaging and quantification of the structural elements of blood clots using high-resolution SEM. (**A**) Selected portions of scanning electron micrographs of clots, illustrating the structures analyzed in this study: a biconcave RBC (*1*), intermediate mainly biconcave RBCs (*2*), intermediate mainly polyhedral RBCs (*3*), a polyhedral compressed RBC (polyhedrocyte) (*4*), fibrin (*5*). Magnification bar = 5 μm. (**B**) A scanning electron micrograph with overlaid grid (the size of each square is 6 μm x 6 μm) used to quantify the composition of a clot. Each grid square contains several cells and/or fibrin that were marked and measured. The relative number for each RBC type and relative fibrin/pores area per image were counted. Magnification bar = 6 μm.

## Results

### Contraction of whole blood and PRP clots of various shapes

The course of clot contraction was reflected by the kinetic curves, where the growing extent of clot contraction was plotted as a function of time tracked over 1 hour (Fig. 4). To reveal the effects of clot shape on contraction dynamics, clots of the same initial volume (0.3 ml or 1.5 ml) were formed and allowed to contract in flat, cuboid, and cylindrical chambers, as shown schematically in Fig. 1. The results presented in Fig. 4A, C and Tables S1, S2 clearly show that the rates and final extents of contraction of whole blood clots were significantly different in the following order: flat > cuboid = cylindrical. For 0.3-ml whole blood clots, a pairwise comparison of the final extent of 1-hour contraction revealed that the median value in the flat clots (80%) was significantly larger compared to the cylindrical blood clots (73%, p=0.006), while the cylindrical (73%) and cuboid (75%) clots had almost the same final extent of contraction (p=0.3). Quantitative comparison of the rates of contraction between the blood clots of various shapes showed a similar dependence on the clot geometry, with a significant difference between the kinetic curves for contraction of flat, cuboid, and cylindrical clots (p<0.0001, Friedman test). For 1.5-ml-clots, only the rates of contraction were significantly different (p<0.0001), but the final extents of clot shrinkage were similar (p=0.8) (Fig. 4C, Table S2).

**Figure 4.**
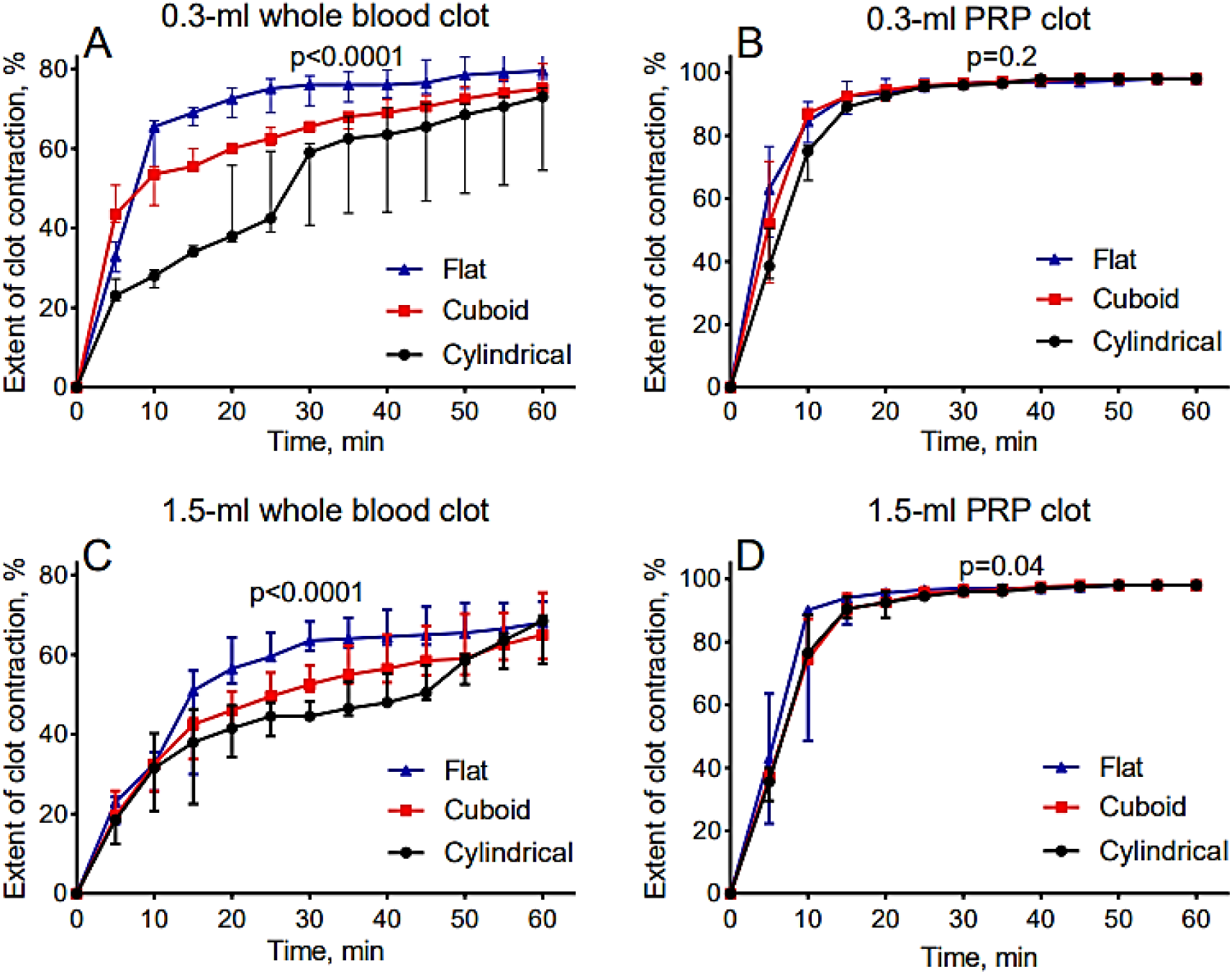
The extent of clot contraction as a function of time for clots of a fixed initial volume but various shapes (flat, cuboid, or cylindrical). (**A**) 0.3-ml whole blood clots of the 3 different shapes (n=3); (**B**) 0.3-ml PRP clots of the 3 different shapes (n=3): (**C**) 1.5-ml whole blood clots of the 3 different shapes (n=3); (**D**) 1.5-ml PRP clots of the 3 different shapes (n=3). The overall p-values determined with the Friedman test reflect the significant difference between the entire kinetic curves (rates) for clots formed both in whole blood and PRP. To assess differences between the extents of contraction at a particular time point, one-way ANOVA test with Holm-Sidak’s multiple comparisons test or Kruskal-Wallis test with Dunn’s multiple comparisons test were used.

Unlike in whole blood clots, in 0.3-ml PRP clots there were no differences observed in the rates of contraction between the flat, cuboid, and cylindrical clots (p=0.2, Friedman test). Pairwise comparisons of the final extent of contraction also revealed that the median values in the cuboid or cylindrical clots (98% each) were equal to the flat blood clots (98%; p=0.2, both) (Fig. 4B, D; Tables S1 and S2). For 1.5-ml PRP clots, there was a moderate difference in the initial contraction rates (p=0.04) that equalized after 20 min and resulted in the same final extent of contraction (Fig. 4D). Apparently, the observed dependence of clot contraction on clot shape was strikingly more pronounced in whole blood than in PRP, suggesting a greatly important role of RBCs as the determinant of shape-dependent variations in blood clot contraction.

Thus, the rate and final extant of contraction of whole blood clots depended on their shape, while in PRP clots the parameters of clot shrinkage were independent of the clot shape at least for the flat, cylindrical, and cuboid clots studied.

### Contraction of whole blood and PRP clots of the same shapes but various dimensions

Next, we analyzed the effects of initial clot size on the rate and extent of clot contraction. The results obtained clearly show that the contraction of whole blood clots of the same shape (cylindrical, cuboid, or flat) varied depending on their initial volume, such that the smaller (0.3 ml) clots always contracted faster and to a larger extent than the larger (1.5 ml) clots (Fig. 5A, C, E; Tables S1-S5). In particular, the final extent of contraction of the smaller 0.3-ml cuboid clots (75%) was larger than 1.5-ml clots (65%, p=0.05) and the smaller 0.3-ml flat clots (80%) were consistently more shrunken than the larger 1.5-ml blood clots (68%, p=0.002). Despite the lack of pairwise statistical significance, the same trend was true for the smaller 0.3-ml cylindrical clots with a median final extent of contraction of 73% compared to the larger 1.5-ml blood clots, which had a 69% median final extent of contraction (p=0.4) (Tables S3-S5).

**Figure 5.**
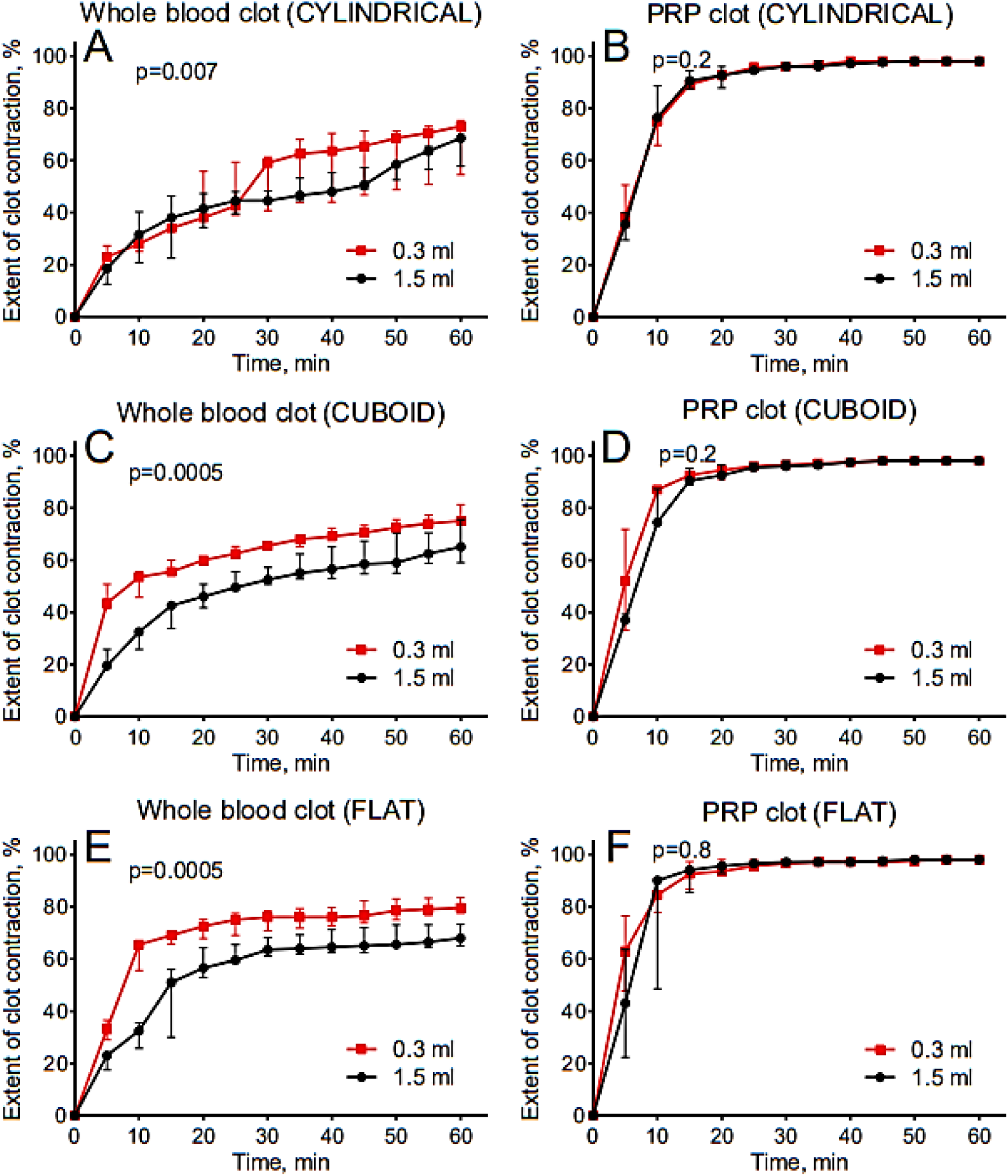
The extent of clot contraction as a function of time for clots of the same shapes (cylindrical, cuboid, or flat) but various dimensions (0.3 ml and 1.5 ml initial volumes). (**A**) Cylindrical whole blood clots; (**B**) Cylindrical PRP clots; (**C**) Cuboid whole blood clots; (**D**) Cuboid PRP clots; (**E**) Flat whole blood clots; (**F**) Flat PRP clots. The p-values determined with Wilcoxon matched-pairs signed rank test reflect the difference between the entire kinetic curves (rates) for clots formed both in whole blood and PRP. To reveal differences between the extents of contraction at each time point, Mann-Whitney test or unpaired *t*-test were used.

In PRP clots, unlike clots in whole blood, there was no discernible difference in the rate and extent of contraction between cylindrical, cuboid, and flat clots of various initial volumes studied (Fig. 5B, D, F). In all PRP clots, there was no difference between 0.3-ml and 1.5-ml clots in the final extent of contraction, but for the cylindrical and cuboid clots there was a moderate stochastic deviation towards the higher (cuboid) or lower (cylindrical) extent of contraction in the smaller 0.3-ml clots at the 10-min time point (Fig. 5B, D; Tables S3-S5).

These distinctions between whole blood and PRP clots again suggest an important role of RBCs in blood clot biomechanics. It is noteworthy that regardless of the initial clot volume (0.3 ml or 1.5 ml), the rate and extent of contraction were always higher in PRP clots compared to whole blood clots, due to the large volume fraction (hematocrit) occupied by RBCs (Fig. 5). Because the observed shape- and size-dependence of clot contraction was substantially more prominent in physiologically relevant whole blood clots, PRP clots were excluded from further structural analysis of contracted clots of various shapes and sizes.

### Contraction of cylindrical blood clots with various dimensions

To mimic the contraction of elongated thrombi formed in blood vessels of various diameters and lengths, we created cylindrical whole blood clots of different dimensions. Cylindrical thrombi can be attached to the vessel wall but they often have a part that floats freely in the vessel lumen; our model reflects mostly the elongated floating part of a thrombus, which is usually larger than the attached portion, which may be flattened during contraction [42]. We sought to reveal if the rate and degree of shrinkage of cylindrical clots depend on their dimensions, which determine variations in the clot volume.

First, we formed cylindrical blood clots with **a constant initial volume (1.5 ml) but distinct diameters and lengths**. The initial clot diameter/length values were as follows: 6.2 mm/50 mm, 9.1 mm/23 mm, and 13.8 mm/10 mm (Fig. 6A). Among the three varieties, the narrowest/longest clots demonstrated the highest rate and final extent of contraction. Specifically, blood clots with the smallest 6.2-mm diameter and the largest 50-mm length (with a median final extent of contraction 63%) shrank more than clots with an intermediate 9.1-mm diameter and the 23-mm length (43%; p=0.01) and the largest 13.8-mm diameter and the shortest 10-mm length (42%; p=0.0006), with the latter two values being similar (Fig. 6A, Table S6). Therefore, the overall order of the extents of clot contraction was as follows: small diameter/large length > intermediate diameter/intermediate length = large diameter/short length.

**Figure 6.**
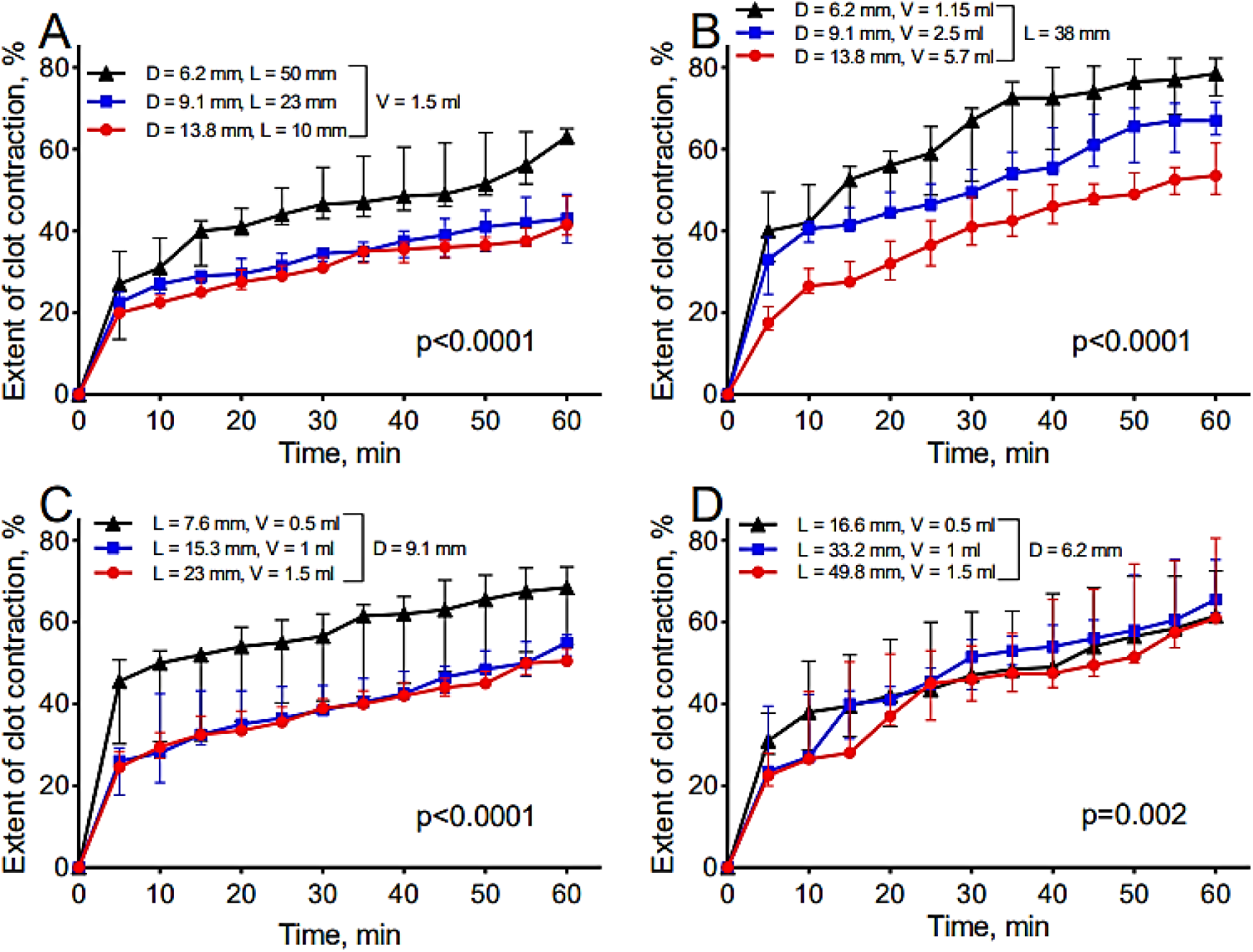
The extent of clot contraction as a function of time for cylindrical clots of various diameters, lengths, and initial volumes formed in whole blood. (**A**) Various diameters (*D*) and corresponding lengths (*L*) but a constant initial volume (*V=1.5 ml*); (**B**) various diameters (*D*) and corresponding initial volumes (*V*) but a constant length (*L=38 mm*); (**C**) various lengths (*L*) and corresponding initial volumes (*V*) but a constant diameter (*D=9.1 mm*); (**D**) various lengths (*L*) and corresponding initial volumes (*V*) but a constant diameter (*D=6.2 mm*) (n=3 for each condition). The p-values determined with Friedman test reflect a difference between the entire kinetic curves (rates) of clot contraction.

Next, we created cylindrical blood clots with **a constant length (38 mm) and various diameters corresponding to distinct initial clot volumes**. The diameter/volume values were: 6.2 mm/1.15 ml, 9.1 mm/2.5 ml, and 13.8 mm/5.7 ml (Fig. 6B). In these clots, the rates and final extents of contraction followed the order: small diameter/small volume > intermediate diameter/intermediate volume > large diameter/large volume. In particular, the smallest clots (6.2 mm length/1.15 ml volume) showed the highest final extent of contraction (79%), which was significantly different from that of the largest clots (13.8 mm length/5.7 ml volume, 54%; p=0.0004), while pairwise comparisons of both with the intermediate clots (67%) showed no significant differences (p>0.05) (Fig. 6B, Table S6).

Finally, we produced cylindrical blood clots with **a constant diameter (9.1 mm or 6.2 mm) and various lengths and initial volumes**. For the clots with a 9.1-mm diameter, the length/volume values were as follows: 23 mm/1.5 ml, 15.3 mm/1 ml, and 7.6 mm/0.5 ml (Fig. 6C, Table S7) and for the clots with a 6.2-mm diameter the length/volume values were as follows: 49.8 mm/1.5 ml, 33.2 mm/1 ml, and 16.6 mm/0.5 ml (Fig. 6D, Table S7). For the clots with a **9.1-mm** diameter, the rates and final extents of contraction followed the following order of clot lengths and initial volumes: small volume/large length > intermediate volume/intermediate length ≥ large volume/small length. In contrast, the extents of contraction were indistinguishable in cylindrical blood clots with a constant diameter **6.2 mm** and various lengths and initial volumes (Fig. 6D and Table S7).

The results indicate that all three variable parameters studied (namely, a cylindrical clot volume, length, and diameter) influence the rate and final extent of blood clot contraction. When the initial clot volume is fixed, clots with a smaller diameter and larger length exhibit stronger contraction. With a constant length, clots with a smaller volume and smaller diameter contract more effectively. When the diameter is constant, clots with a smaller volume and larger length undergo a somewhat faster contraction. Overall, the initial volume, diameter, and length each have distinct impacts on the kinetics and extent of cylindrical clot contraction. In particular, the diameter is a more important determinant of clot contraction, while the clot length is less significant. In other words, the extent of contraction of a cylindrical clot or thrombus depends strongly on its thickness rather than on its length.

### Composition and structural non-uniformity of contracted cylindrical blood clots of various initial dimensions

Given that the elongated clot shape mimics the most common intravascular thrombi, cylindrical blood clots were subjected to ultrastructural examination after 1-hour of contraction. To study meticulously and compare the composition of fully contracted cylindrical blood clots with distinct initial volumes (0.5 ml and 1.5 ml), high-resolution SEM was employed. The main structural components of blood clots are depicted in Fig. 3, and their relative content throughout the clots is presented in Fig. 7A, C, E and Table S8. To analyze the spatial segregation of clot components, images of transversely sectioned contracted clots were visually divided into three layers characterized by distinct packing densities: a looser clot periphery, moderately compact intermediate layer, and tightly packed central portion (Fig. 2). The composition of each of the three layers (peripheral, intermediate, and central) was separately analyzed for both the smaller (0.5 ml) and larger (1.5 ml) contracted cylindrical blood clots (Fig. 7B, D, F; Table S8).

**Figure 7.**
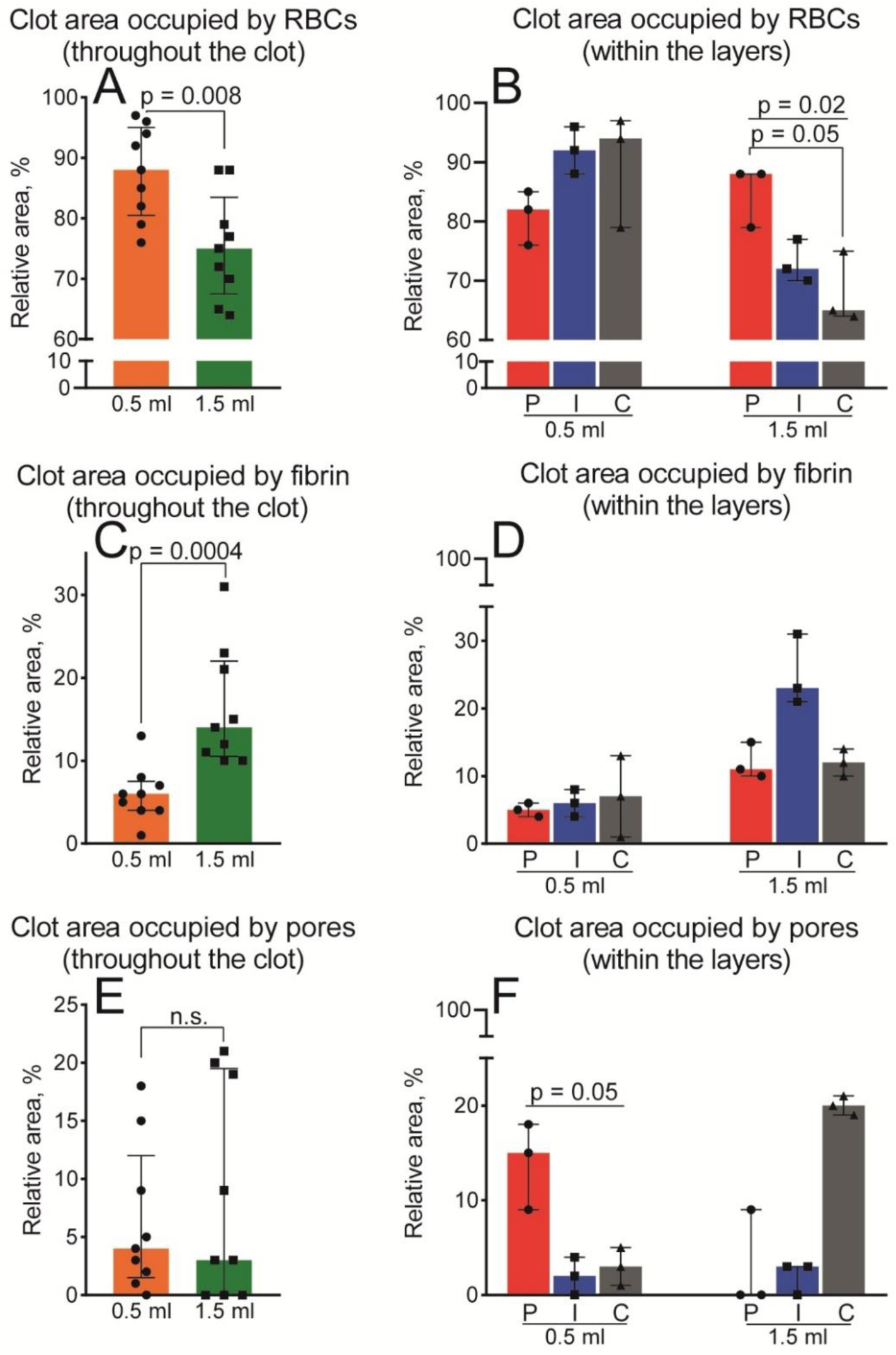
Relative content and spatial distribution of the structural elements of the contracted cylindrical clots with a 0.5-ml versus 1.5-ml initial volume as visualized with SEM. (**A, C, E**) Average relative area occupied by RBCs (A), fibrin (C), and pores (E) throughout the smaller and larger blood clots. Each dot represents the number obtained from an individual SEM image (n=9; 3 images from each of the 3 layers). **(B, D, F)** Relative area occupied by RBCs (B), fibrin (D), and pores (F) within the peripheral (*P*), intermediate (*I*), and central (*C*) layers of the smaller and larger contracted cylindrical blood clots. Each dot represents the number obtained from an individual SEM image (n=3). Results are presented as the median (a top edge of the column) and interquartile range (error bars). Statistical analysis: Kruskal-Wallis test with Dunn’s multiple comparisons test (B, D, F) and Mann-Whitney test (A, C, E). Designation: n.s. – not significant (p>0.05).

The most abundant component of the contracted clots was **RBCs**, comprising approximately 80-90% of the clot volume (Fig. 7A). Remarkably, in the smaller 0.5-ml clots, RBCs occupied a larger relative bulk space (median relative content 88%) than in the 1.5-ml clots (75%; p=0.008) (Table S8), likely due to expulsion of serum and reduction of the intercellular space. In the smaller clots (0.5 ml), there was a clear trend towards increasing RBC accumulation in the central (94%) and intermediate (92%) parts compared to the periphery (82%; p=0.2), which is a known feature of clot contraction [1]. Conversely, in the larger clots (1.5 ml), RBCs accumulated predominantly in the peripheral part (88%) compared to the intermediate layer (72%; p=0.02) and the center of the clot (65%; p=0.05) (Fig. 7B).

In contrast to RBCs, the relative bulk area occupied by **fibrin** was significantly greater in the larger 1.5-ml clots (14%) compared to the smaller 0.5-ml clots (6%; p=0.0004) (Fig. 7C, Table S8), because a lower extent of contraction is associated with less compaction and densification of the fibrin network, which is a characteristic structural feature of a fully contracted clot [43]. In the smaller cylindrical clots (0.5 ml), fibrin was evenly distributed without a significant difference between the layers (p=0.8). Similarly, the distribution of fibrin within the layers of the larger clots did not reveal any substantial difference (p=0.07). However, despite the lack of statistical significance, fibrin appeared to predominate in the intermediate part of the larger clot (Fig. 7D).

Although the **average porosity** measured as the area occupied by empty spaces was similar in the larger and smaller contracted clots (4% and 3%, respectively, p=0.9), the porosity was spatially non-uniform with a higher porosity observed in the middle of the larger clot (Fig. 7E; Table S8). The porosity of the smaller clots was prevalent in the peripheral part (median 15%) versus the intermediate layer (2%) and central portion (3%) (p=0.05, Kruskal-Wallis test). Porosity in the central part of the larger clot (20%) visually dominated over the intermediate layer (3%) and clot periphery (zero porosity); however, this apparent overall trend was at the border of statistical significance, due to scattered values (p=0.06) (Fig. 7F).

Thus, in the smaller contracted cylindrical clots (0.5 ml), the increasing accumulation of RBCs occurred in the direction from the periphery to the center, which was inversely related to the porosity, which increased from the center to periphery. In contrast, in the larger clots (1.5 ml), RBCs accumulated in the direction from the center to the periphery, while the porosity increased from the periphery to the center (Fig. 7B, F). Fibrin was more abundant in the larger clots (Fig. 7C), reflecting the lower extent of clot contraction and reduced compaction and densification of fibrin, but it did not have a significantly distinct distribution between the layers in the smaller versus larger clots (Fig. 7D).

### Overall content and spatial non-uniformity of RBCs with various degrees of compression in the smaller and larger cylindrical blood clots

The high-resolution SEM enabled us to assess not only the overall RBC content and gradients but also the transition from customary biconcave RBCs to compressed polyhedral RBCs, which comprises an important morphological feature of contracted blood clots both *in vitro* and *in vivo* [*1*; *8; 9; 14-18*]. The identified RBC types analyzed in the contracted clots included fully compressed polyhedral, intermediately compressed mainly polyhedral, slightly compressed intermediate mainly biconcave, and uncompressed native biconcave RBCs (see Methods and Fig. 3).

First, to assess the degree of bulk clot compression, we quantified the average content of non-deformed and deformed RBCs throughout the cylindrical clots of different initial sizes, 0.5 ml and 1.5 ml (Fig. 8A, C, E, G; Table S9). The overall content of compressed RBCs was much higher in the smaller 0.5-ml clots (median fractions of RBCs: 8% polyhedral and 41% mainly polyhedral), while in the larger 1.5-ml clots their content was substantially smaller with zero median numbers for polyhedral and mainly polyhedral RBCs (p=0.0006) (Fig. 8A, C). On the contrary, uncompressed biconcave and only slightly compressed mainly biconcave RBCs predominated in the larger 1.5-ml clots (median 5% and 85%, respectively), in comparison with 0.5-ml clots that contained zero biconcave and only 14% mainly biconcave RBCs (p=0.004) (Fig. 8E, G). These differences in the fractions of uncompressed and compressed RBCs confirm morphologically that the bulk compaction of the smaller clots was substantially more evident than in the larger clots (Table S9).

**Figure 8.**
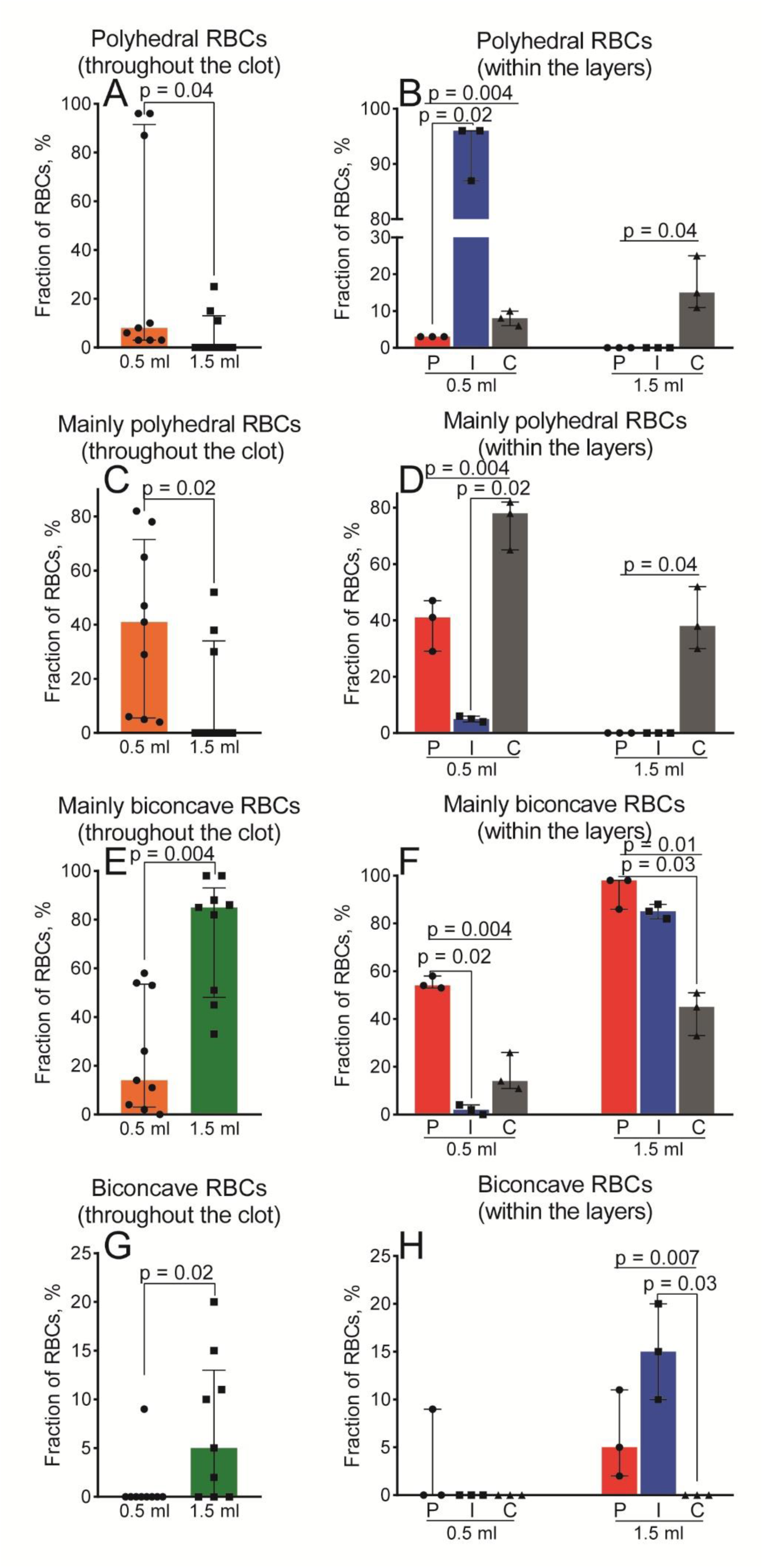
Relative fractions of RBCs in contracted cylindrical clots having a 0.5-ml versus 1.5-ml initial volume as visualized with SEM. **(A, C, E, G)** Averaged fractions of polyhedral RBCs (A), intermediate mainly polyhedral RBCs (C), intermediate mainly biconcave RBCs (E) and biconcave RBCs (G) throughout the clots. Each dot represents the fraction of the RBC types obtained from individual SEM images of cylindrical blood clots (n=9; 3 images from each of 3 the layers). **(B, D, F, H)** Fractions of polyhedral RBCs (B), intermediate mainly polyhedral RBCs (D), intermediate mainly biconcave RBCs (F) and biconcave RBCs (H) within the peripheral (*P*), intermediate (*I*), and central (*C*) layers of fully contracted cylindrical blood clots. Each dot represents a fraction of a certain RBC type obtained from individual SEM images of cylindrical blood clots (n=3). Results are presented as the median (a top edge of the column) and interquartile range (error bars). Statistical analysis: Kruskal-Wallis test with Dunn’s multiple comparisons test (B, D, F, H) and Mann-Whitney test (A, C, E, G). In multiple comparisons, the p-values show significant differences within one subgroup (0.5 ml or 1.5 ml), and not between them, in order to show the gradient of cell redistribution within a clot of the same volume.

To assess the spatial non-uniformity of blood clot contraction and its dependence on clot size, the RBC fractions were quantified in each of the three layers (peripheral, intermediate, and central), discerned based on their packing density (see Fig. 2).

The results indicate that in the smaller 0.5-ml cylindrical clots, compressed RBCs (polyhedral and mainly polyhedral) were present in all three layers of the clot with a moderate increase from the periphery through the center. Conversely, in the larger 1.5-ml clots, polyhedral and mainly polyhedral RBCs were only found in the central part of the contracted clot (Fig. 8B, D; Table S9). Despite the common trend towards centralization, the median content of compressed RBCs in the center of the larger clot was only 53% (15% polyhedral plus 38% mainly polyhedral), while in the center of the smaller clots, the combined content of compressed RBCs was 83% (8% polyhedral plus 78% mainly polyhedral, p=0.45) (Fig. 8B, D). Accordingly, in the central part of the smaller clots, the relative count of only slightly compressed mainly biconcave RBCs (14%) was about three times lower than in the larger clots (45%) (p=0.02) (Fig. 8F; Table S9), indicating the biggest compression of RBCs in the center of the smaller clots.

Remarkably, the intermediate layer of the smaller clots was characterized by a significant predominance of polyhedral RBCs (median 96%), indicating that the strongest contractile stress was produced in the intermediate part of the smaller clots. In contrast, the intermediate layer of the larger clots contained mostly slightly compressed mainly biconcave cells (85%) and uncompressed biconcave (15%) RBCs without any fully compressed polyhedrocytes, indicating extremely weak compression of the intermediate part of the larger clots (Fig. 8B, D, F, H; Table S9).

The peripheral part of the smaller clots contained partially compressed cells, namely mainly polyhedral (41%) and mainly biconcave (54%) RBCs (median total ∼95%), with a small content of polyhedral RBCs (3%) (Fig. 8B, D; Table S9). In contrast, in the peripheral layer of the larger clots, only slightly compressed RBCs were present, namely mainly biconcave cells (98%) without polyhedral cells at all, and a small fraction of uncompressed biconcave forms (5%) (Fig. 8F, H; Table S9).

These data indicate distinct spatial distributions of compressive stresses within the smaller and larger cylindrical clots, leading to dissimilar spatial spreading of RBCs with various degrees of compression. Specifically, in the larger cylindrical clots (1.5 ml), only the central part undergoes moderate contraction, while in the smaller clots (0.5 ml), much stronger contraction and compression occur throughout the clot, especially in the intermediate layer between the clot periphery and the core.

## Discussion

The present study aims at investigating whether, in addition to the extrinsic modulators, contraction of blood clots and thrombi depends on their inherent fundamental properties, such as geometry and size, which are quite variable *in vivo* and depend on the location of the thrombus and conditions of its formation.

The main finding is that platelet-driven contraction of a blood clot does depend on both its shape and size. To compare distinct though simplified geometries, we analyzed in parallel fresh clots from the same human blood samples made in cuboid, flat, and cylindrical containers that varied in the initial clot size. Of the three types of shape studied, the propensity of clots to contract has the following order: flat > cuboid = cylindrical (Figs. 4A,C; 5A,C,E). The physiological relevance and importance of this finding is that the observed dependence reflects the modulating role on clot contraction of the shape of different parts of the blood circulation system (blood vessels, heart cavities, brain sinuses, wounds, aneurysms, etc.) in which a blood clot or a thrombus can be formed. To be more specific, venous thrombi form gradually under conditions of stasis/reduced blood flow and decreased wall shear stress [39], whereas arterial thrombi form more or less rapidly under conditions of fast blood flow (high shear rates within the stenotic region [44]), with a complex, non-uniform distribution of components. Accordingly, venous thrombi are usually more longitudinal, spatially voluminous and obstructive (replicating the shape of the vessel) than arterial thrombi, which can be parietal and flattened [45–51]. In addition, venous thrombi more resemble *in vitro* clots, in that they contain more RBCs and are more homogeneous in structure [16]. Stenosis of blood vessels associated with disturbed blood flow can also contribute to the shape of thrombi, making them narrower [52–54]. Obviously, the hemodynamic conditions together with the geometry of vessels, in which blood clots form, determine the shape of thrombi. To the best of our knowledge, this aspect of hemostasis and thrombosis related to variations in clot geometries has not been previously addressed.

The size-dependence of blood clot contraction is another important and physiologically relevant discovery. In blood clots of all the shapes studied, the smaller clots always contracted faster and to a larger extent than the larger clots (Figs. 4A,C; 5A,C,E). To study this phenomenon in more detail, we used cylindrical clots, as they reproduce the shape of a blood vessel lumen and comprise the most common geometry of obstructive intravascular thrombi. Dimensions of cylindrical clots are characterized by three parameters, namely volume, diameter, and length, and each of those was varied to form clots of various sizes. Our results clearly show that each of the three parameters has impact on the contraction of cylindrical clots, such that a smaller volume/diameter/length resulted in a higher extent of clot contraction (Fig. 6). Notably, the impact of the clot diameter on contraction is greater than the effect of the length. Since a higher extent of contraction makes smaller thrombi less occlusive and more stable, the observed size-dependence has significant clinical implications confirmed indirectly by a number of observations. In particular, thrombus burden (assessed with the Mastora score based on the percentage of embolic cross-sectional area of the pulmonary artery) correlates directly with a larger superior vena cava diameter >20 mm in patients with acute pulmonary embolism [*55; 56*]. The mortality rate in pulmonary embolism was found to be associated with a larger diameter (>29 mm) of the pulmonary artery trunk [57]. The relative clot size and stenotic degree in pulmonary embolism are significant predictors of a perfusion defect on conventional computed tomographic pulmonary angiography, such that lesions with higher degrees of stenosis (>80%) have higher percentages of perfusion defect because of less contraction [58]. Apparently, our finding of the larger clots being less contracted and hence potentially more obstructive is consistent with the greater threat for the larger emboli.

It is noteworthy that blood clots studied here correspond to the dimensions of thrombi formed in the medium (∼1-8 mm in diameter) and large-caliber (>8 mm) vessels, which are often involved in some of the most significant thrombotic conditions. In particular, the normal diameter of coronary arteries measures between 3 and 5 mm [*59; 60*] and a coronary thrombus is considered "small" when its greatest dimension is less than half the diameter of the coronary artery, while a "large" thrombus is defined as having a dimension greater than twice the vessel diameter [61]. Pre-cerebral and cerebral vessel diameters can vary due to patient-specific factors and imaging techniques, but they range roughly between 2 and 5 mm [*62; 63*]. It should be noted that studies of *ex vivo* thrombi from these patient groups have shown that they contain large amounts of both fibrin and RBCs and show evidence of clot contraction [15]. Extracranial segments of the carotid artery have a diameter of 6-9 mm with an age-related increase in the diameter associated with the risk of cardiovascular death [64]. The normal diameter of the abdominal aorta is usually less than 3.0 cm [65] and the risk of complications increases as the aorta grows [66]. These facts support the physiological relevance of our study in a number of respects: i) the reported thrombus sizes correspond to the sizes of the clots we studied; ii) again, the trend reported for the size of a thrombus fits our data indicating that larger clots are less contracted and therefore more obstructive; iii) the shape of most arterial thrombi is cylindrical, which is the major clot shape we studied.

The diameter of veins varies by location; in particular, the inferior vena cava is 1.2-1.7 cm in diameter, iliac veins near the groin are greater than 1 cm, common femoral vein diameter is about 1 cm, and superficial and deep femoral vein diameters are usually 6-7 mm. Notably, vein segments of the lower limbs with acute thrombosis are larger compared to normal veins [67–69]. The size of a venous thrombus depends on the vein in which it occurs. A clot diameter of at least 5 mm is generally considered to be deep vein thrombosis (DVT). However, about a quarter of patients with DVT have a clot diameter <5 mm. A thrombus length of 9 cm or more is likely to indicate thrombus extension and could support a diagnosis of recurrent DVT [*49; 70*]. Notably, the clots studied here resemble the shape and size variations reported for venous thrombi.

Importantly, the dependency of contraction on the shape and size of clots has been found only for whole blood clots, while in PRP clots the dependence is negligibly small or absent (Figs. 4B,D; 5B,D,F), indicating a critically important role of RBCs in clot biomechanics. Although many arterial thrombi contain a substantial proportion of RBCs (e.g. about 17% for coronary artery thrombi) [*15; 16*], the results here will apply more to venous thrombi, which are richer in RBCs and fibrin. RBCs occupy the majority of whole blood clot volume and impede clot contraction as a key mechanically resilient clot component [*19; 71; 72*]. In addition, during clot contraction, RBCs are accumulated in the core of a clot and change their shape from biconcave to polyhedral with a simultaneous increase in cell packing density [1]. In the interior of a contracted blood clot, both electron and light microscopy reveal compressed RBCs that have an unusual polyhedral shape, named polyhedrocytes or piezocytes [14]. Unlike RBCs, in contracted blood clots and thrombi platelets and fibrin segregate towards the outer layer, and the redistribution of clot components is a main morphological signature of clot contraction in both venous and arterial thrombi [*1; 7*].

These morphological signs of contraction, namely formation of polyhedral RBCs and spatial redistribution of clot components, have been analyzed thoroughly in the smaller versus larger cylindrical clots, corresponding to the real blood clots observed in blood vessels *in vivo* [36]. The results obtained have led us to a number of important conclusions. 1) In the smaller contracted clots, RBCs comprise a greater volume fraction than in the larger clots (Fig. 7A), which is consistent with a higher extent of contraction, implying more serum expulsion and reduction in the intercellular space. 2) The smaller contracted clots have a higher content of polyhedrocytes, along with distinctions in the distribution of compressed RBCs between clot layers (Figs. 8A-D). This finding is in line with the observation that the overall bulk compaction of the smaller clots is substantially more pronounced than in the larger clots (Figs. 4-8). 3) In the smaller clots, an increase in the accumulation of RBCs and a decrease in porosity occurred in the direction from the periphery to the center of the clot (Figs. 7B, F). On the contrary, in the larger clots, an ascending accumulation of RBCs and descending porosity occurred in the direction from the core to the edge of the clot, indicating an atypical spatial redistribution of clot components, which reflects weaker contractile forces hardly penetrating the bulky mass of the larger contracting clots. 4) The distinct spatial non-uniformity of the clot contraction process between the smaller and larger clots has even more morphological signatures. In the larger clots, only the central part undergoes moderate contraction, while in the smaller clots, much stronger contraction and cellular compression occurred throughout the clot, especially in the intermediate layer (Fig. 8B), showing distinct spatial distribution of compressive stresses within the smaller and larger cylindrical clots and/or different amount of serum expelled from larger versus smaller contracted clots. Moreover, the spatial spreading of clot components between the clot periphery and center in the larger clots is mostly opposite to that of the smaller cylindrical contracted blood clots (Fig. 7B, F). Altogether, the structural data indicate that in the larger clots, contraction is less and spatially more heterogeneous, likely because the contractile force of activated platelets is relatively insufficient to effectively compress through a bulky, large mass clot with a higher overall mechanical resilience and/or likely because of lesser serum expulsion.

The results obtained in this study have great pathophysiological importance because blood clot size and geometry may affect the vitally important biological and mechanical thrombus properties determined by clot contraction. It has been known that clot contraction enhances the rate of internal fibrinolysis ∼2-fold [10]. Accordingly, in the smaller clots the efficacy of internal fibrinolysis should be greater, resulting in reduction of thrombus size, lower obstructiveness, and reduced clot durability [73]. On the other hand, external fibrinolysis is ∼4-fold slower in contracted clots [10], which potentially makes the smaller clots less susceptible to therapeutic thrombolysis. The relationships between thrombus size and susceptibility to mechanical stability may be more complicated. Since the bulky thrombi are less contracted/compacted and softer, they may be more susceptible to mechanical thrombectomy than the smaller and stiffer thrombi. On the other hand, the bulky intravital thrombi can be more prone to rupture and embolization, which has been shown for the younger and less contracted portions of venous thrombi [9]. Based on our data, the thrombi larger than 1-1.5 ml are less contracted and their structure and composition is highly heterogeneous and anisotropic, making the progression of thrombosis less predictable. Anyway, the shape and size of a thrombus should be considered clinically important parameters, affecting the course and outcome of thrombosis.

In summary, this study has revealed that the initial shape and volume of a blood clot both affect the rate and extent of platelet-driven clot contraction. Flat clots contract more than cuboid or cylindrical clots and the smaller clots shrink more and faster, whereas contraction of larger clots is slower and less complete, with a high degree of structural non-uniformity. Unlike clots formed with whole blood, plasma clots display no dependence of contraction on clot shapes and volumes, suggesting a key role of RBCs in this aspect of blood clot mechanics. These novel findings are pathophysiologically and clinically relevant as they provide insights on the role of variable geometry and size of intravascular thrombi in mechanical remodeling that largely determines their biological and physical properties.

## Acknowledgements

Scanning electron microscopy was carried out in the Interdisciplinary Center for Analytical Microscopy of Kazan Federal University.

## Sources of Funding

The work was supported by the NIH grants PO1-HL146373 and RO1-148227.

## Abbreviations and Acronyms

RBC: red blood cell
ATP: adenosine triphosphate
PRP: platelet-rich plasma
SEM: scanning electron microscopy
ANOVA: analysis of variance
CT: computed tomography
DVT: deep vein thrombosis

## Disclosures

None.

## Novelty and Significance

### What is known?

- Intravascular blood clots and thrombi are characterized by the variable geometry/size.
- Blood clots undergo the volumetric shrinkage (contraction).

### What new information does this article contribute?

- The smaller and larger clots have distinct size-dependent rates and extents of platelet-driven contraction as well as distinct degrees of structural/spatial non-uniformity, reflecting different spatial gradients of compressive stresses.
- The platelet-rich plasma clots contract independently of the clot dimensions and shapes, indicating a key role of erythrocytes.
- The patophysiological and clinical relevance is related to the variable geometry/size and their impact on the outcomes of intravascular thrombosis.

## Notes

### Competing Interest Statement

The authors have declared no competing interest.

## References

1. Cines DB, Lebedeva T, Nagaswami C, Hayes V, Massefski W, Litvinov RI, Rauova L, Lowery TJ, Weisel JW. Clot contraction: compression of erythrocytes into tightly packed polyhedra and redistribution of platelets and fibrin. Blood. 2014;123:1596–1603. doi: 10.1182/blood-2013-08-523860

2. Stalker TJ, Welsh JD, Tomaiuolo M, Wu J, Colace TV, Diamond SL, Brass LF. A systems approach to hemostasis: 3. Thrombus consolidation regulates intrathrombus solute transport and local thrombin activity. Blood. 2014;124:1824–1831. doi: 10.1182/blood-2014-01-550319

3. Lam WA, Chaudhuri O, Crow A, Webster KD, Li TD, Kita A, Huang J, Fletcher DA. Mechanics and contraction dynamics of single platelets and implications for clot stiffening. Nat Mater. 2011;10:61–66. doi: 10.1038/nmat2903

4. Leong L, Chernysh IN, Xu Y, Sim D, Nagaswami C, de Lange Z, Kosolapova S, Cuker A, Kauser K, Weisel JW. Clot stability as a determinant of effective factor VIII replacement in hemophilia A. Res Pract Thromb Haemost. 2017;1:231–241. doi: 10.1002/rth2.12034

5. Nurden AT. Molecular basis of clot retraction and its role in wound healing. Thromb Res. 2023;231:159–169. doi: 10.1016/j.thromres.2022.08.010

6. Tutwiler V, Wang H, Litvinov RI, Weisel JW, Shenoy VB. Interplay of platelet contractility and elasticity of fibrin/erythrocytes in blood clot retraction. Biophys J. 2017;112:714–723. doi: 10.1016/j.bpj.2017.01.005

7. Litvinov RI, Weisel JW. Blood clot contraction: mechanisms, pathophysiology, and disease. Res Pract Thromb Haemost. 2022;7:100023. doi: 10.1016/j.rpth.2022.100023

8. Peshkova AD, Malyasyov DV, Bredikhin RA, Le Minh G, Andrianova IA, Tutwiler V, Nagaswami C, Weisel JW, Litvinov RI. Reduced contraction of blood clots in venous thromboembolism is a potential thrombogenic and embologenic mechanism. TH Open. 2018;2:e104–e115. doi: 10.1055/s-0038-1635572

9. Khismatullin RR, Abdullayeva S, Peshkova AD, Sounbuli K, Evtugina NG, Litvinov RI, Weisel JW. Extent of intravital contraction of arterial and venous thrombi and pulmonary emboli. Blood Adv. 2022;6:1708–1718. doi: 10.1182/bloodadvances.2021005801

10. Tutwiler V, Peshkova AD, Le Minh G, Zaitsev S, Litvinov RI, Cines DB, Weisel JW. Blood clot contraction differentially modulates internal and external fibrinolysis. J Thromb Haemost. 2019;17:361–370. doi: 10.1111/jth.14370

11. Carr ME Jr, Zekert SL. Measurement of platelet-mediated force development during plasma clot formation. Am J Med Sci. 1991;302:13–18. doi: 10.1097/00000441-199107000-00004

12. Jen CJ, McIntire LV. The structural properties and contractile force of a clot. Cell Motil. 1982;2:445–455. doi: 10.1002/cm.970020504

13. Qiu Y, Ciciliano J, Myers DR, Tran R, Lam WA. Platelets and physics: how platelets "feel" and respond to their mechanical microenvironment. Blood Rev. 2015;29:377–386. doi: 10.1016/j.blre.2015.05.002

14. Tutwiler V, Mukhitov AR, Peshkova AD, Le Minh G, Khismatullin RR, Vicksman J, Nagaswami C, Litvinov RI, Weisel JW. Shape changes of erythrocytes during blood clot contraction and the structure of polyhedrocytes. Sci Rep. 2018;8:17907. doi: 10.1038/s41598-018-35849-8

15. Khismatullin RR, Nagaswami C, Shakirova AZ, Vrtková A, Procházka V, Gumulec J, Mačák J, Litvinov RI, Weisel JW. Quantitative morphology of cerebral thrombi related to intravital contraction and clinical features of ischemic stroke. Stroke. 2020;51:3640–3650. doi: 10.1161/STROKEAHA.120.031559

16. Chernysh IN, Nagaswami C, Kosolapova S, Peshkova AD, Cuker A, Cines DB, Cambor CL, Litvinov RI, Weisel JW. The distinctive structure and composition of arterial and venous thrombi and pulmonary emboli. Sci Rep. 2020;10:5112. doi: 10.1038/s41598-020-59526-x

17. Litvinov RI, Khismatullin RR, Shakirova AZ, Litvinov TR, Nagaswami C, Peshkova AD, Weisel JW. Morphological signs of intravital contraction (retraction) of pulmonary thrombotic emboli. BioNanoScience. 2018;8:428–433. doi: 10.1007/s12668-017-0476-1

18. Ząbczyk M, Sadowski M, Zalewski J, Undas A. Polyhedrocytes in intracoronary thrombi from patients with ST-elevation myocardial infarction. Int J Cardiol. 2015;179:186–187. doi: 10.1016/j.ijcard.2014.10.004

19. Tutwiler V, Litvinov RI, Lozhkin AP, Peshkova AD, Lebedeva T, Ataullakhanov FI, Spiller KL, Cines DB, Weisel JW. Kinetics and mechanics of clot contraction are governed by the molecular and cellular composition of the blood. Blood. 2016;127:149–159. doi: 10.1182/blood-2015-05-647560

20. Safiullina SI, Evtugina NG, Andrianova IA, Khismatullin RR, Kravtsova OA, Khabirova AI, Nagaswami C, Daminova AG, Peshkova AD, Litvinov RI, et al. A familial case of MYH9 gene mutation associated with multiple functional and structural platelet abnormalities. Sci Rep. 2022;12:19975. doi: 10.1038/s41598-022-24098-5

21. Sun Y, Myers DR, Nikolov SV, Oshinowo O, Baek J, Bowie SM, Lambert TP, Woods E, Sakurai Y, Lam WA, et al. Platelet heterogeneity enhances blood clot volumetric contraction: An example of asynchrono-mechanical amplification. Biomaterials. 2021;274:120828. doi: 10.1016/j.biomaterials.2021.120828

22. Carr ME Jr, Martin EJ, Carr SL. Delayed, reduced or inhibited thrombin production reduces platelet contractile force and results in weaker clot formation. Blood Coagul Fibrinolysis. 2002;13:193–197. doi: 10.1097/00001721-200204000-00004

23. Carr ME Jr. Development of platelet contractile force as a research and clinical measure of platelet function. Cell Biochem Biophys. 2003;38:55–78. doi: 10.1385/CBB:38:1:55

24. Peshkova AD, Le Minh G, Tutwiler V, Andrianova IA, Weisel JW, Litvinov RI. Activated monocytes enhance platelet-driven contraction of blood clots via tissue factor expression. Sci Rep. 2017;7:5149. doi: 10.1038/s41598-017-05601-9

25. Aleman MM, Byrnes JR, Wang JG, Tran R, Lam WA, Di Paola J, Mackman N, Degen JL, Flick MJ, Wolberg AS. Factor XIII activity mediates red blood cell retention in venous thrombi. J Clin Invest. 2014;124:3590–3600. doi: 10.1172/JCI75386

26. Byrnes JR, Duval C, Wang Y, Hansen CE, Ahn B, Mooberry MJ, Clark MA, Johnsen JM, Lord ST, Lam WA, et al. Factor XIIIa-dependent retention of red blood cells in clots is mediated by fibrin α-chain crosslinking. Blood. 2015;126:1940–1948. doi: 10.1182/blood-2015-06-652263

27. Tutwiler V, Peshkova AD, Andrianova IA, Khasanova DR, Weisel JW, Litvinov RI. Contraction of blood clots is impaired in acute ischemic stroke. Arterioscler Thromb Vasc Biol. 2017;37:271–279. doi: 10.1161/ATVBAHA.116.308622

28. Evtugina NG, Peshkova AD, Pichugin AA, Weisel JW, Litvinov RI. Impaired contraction of blood clots precedes and predicts postoperative venous thromboembolism. Sci Rep. 2020;10:18261. doi: 10.1038/s41598-020-75234-y

29. Peshkova AD, Safiullina SI, Evtugina NG, Baras YS, Ataullakhanov FI, Weisel JW, Litvinov RI. Premorbid hemostasis in women with a history of pregnancy loss. Thromb Haemost. 2019;119:1994–2004. doi: 10.1055/s-0039-1696972

30. Le Minh G, Peshkova AD, Andrianova IA, Sibgatullin TB, Maksudova AN, Weisel JW, Litvinov RI. Impaired contraction of blood clots as a novel prothrombotic mechanism in systemic lupus erythematosus. Clin Sci (Lond). 2018;132:243–254. doi: 10.1042/CS20171510

31. Peshkova AD, Evdokimova TA, Sibgatullin TB, Ataullakhanov FI, Litvinov RI, Weisel JW. Accelerated spatial fibrin growth and impaired contraction of blood clots in patients with rheumatoid arthritis. Int J Mol Sci. 2020;21:9434. doi: 10.3390/ijms21249434

32. Tomasiak-Lozowska MM, Rusak T, Misztal T, Bodzenta-Lukaszyk A, Tomasiak M. Reduced clot retraction rate and altered platelet energy production in patients with asthma. J Asthma. 2016;53:589–598. doi: 10.3109/02770903.2015.1130151

33. Andrianova IA, Khabirova AI, Ponomareva AA, Peshkova AD, Evtugina NG, Le Minh G, Sibgatullin TB, Weisel JW, Litvinov RI. Chronic immune platelet activation is followed by platelet refractoriness and impaired contractility. Int J Mol Sci. 2022;23:7336. doi: 10.3390/ijms23137336

34. Vulliamy P, Armstrong PC. Platelets in hemostasis, thrombosis, and inflammation after major trauma. Arterioscler Thromb Vasc Biol. 2024;44:545–557. doi: 10.1161/ATVBAHA.123.318801

35. Guenego A, Fahed R, Sussman ES, Leipzig M, Albers GW, Martin BW, Marcellus DG, Kuraitis G, Marks MP, Lansberg MG, et al. Impact of clot shape on successful m1 endovascular reperfusion. Front Neurol. 2021;12:642877. doi: 10.3389/fneur.2021.642877

36. Misumi K, Hagiwara Y, Ogura T. Giant cylinder-shaped venous thrombus caused by extracorporeal membrane oxygenation cannula. Acute Med Surg. 2022;9:e791. doi: 10.1002/ams2.791

37. Kumar V, Abbas AK, Aster JC. Hemodynamic disorders, thromboembolic disease, and shock. In: Robbins & Cotran Pathologic Basis of Disease, Tenth ed. (International ed). Philadelphia, PA: Elsevier; 2021:115-140. ISBN: 978-0-323-53113-9

38. Levi M. Arterial and venous thrombosis: more in common than previously thought. Neth J Med. 2011;69:3–4.

39. Wolberg AS, Aleman MM, Leiderman K, Machlus KR. Procoagulant activity in hemostasis and thrombosis: Virchow’s triad revisited. Anesth Analg. 2012;114:275–285. doi: 10.1213/ANE.0b013e31823a088c

40. Prandoni P. Venous and arterial thrombosis: is there a link?. Adv Exp Med Biol. 2017;906:273–283. doi: 10.1007/5584_2016_121

41. Löwenberg EC, Meijers JC, Levi M. Platelet-vessel wall interaction in health and disease. Neth J Med. 2010;68:242–251.

42. Yang S, Yu J, Zeng W, Yang L, Teng L, Cui Y, Shi H. Aortic floating thrombus detected by computed tomography angiography incidentally: Five cases and a literature review. J Thorac Cardiovasc Surg. 2017;153:791–803. doi: 10.1016/j.jtcvs.2016.12.015

43. Kim OV, Litvinov RI, Alber MS, Weisel JW. Quantitative structural mechanobiology of platelet-driven blood clot contraction. Nat Commun. 2017;8:1274. doi: 10.1038/s41467-017-00885-x

44. Bark DL Jr, Ku DN. Wall shear over high degree stenoses pertinent to atherothrombosis. J Biomech. 2010;43:2970–2977. doi: 10.1016/j.jbiomech.2010.07.011

45. Gacko M, Worowska A, Głowiński S. Coagulative and fibrinolytic activity in parietal thrombus of aortic aneurysm. Rocz Akad Med Bialymst. 1999;44:102–110

46. Malyar NM, Janosi RA, Brkovic Z, Erbel R. Large mobile thrombus in non-atherosclerotic thoracic aorta as the source of peripheral arterial embolism. Thromb J. 2005;3:19. doi: 10.1186/1477-9560-3-19

47. Drouet L. Atherothrombosis as a systemic disease. Cerebrovasc Dis. 2002;13 Suppl 1:1–6. doi: 10.1159/000047782

48. Panpikoon T, Phattharaprueksa W, Treesit T, Bua-Ngam C, Pichitpichatkul K, Sriprachyakul A. Morphologic change in deep venous thrombosis in the lower extremity after therapeutic anticoagulation. Thromb J. 2021;19:99. doi: 10.1186/s12959-021-00352-0

49. Linkins LA, Pasquale P, Paterson S, Kearon C. Change in thrombus length on venous ultrasound and recurrent deep vein thrombosis. Arch Intern Med. 2004;164:1793–1796. doi: 10.1001/archinte.164.16.1793

50. Kurklinsky AK, Kalsi H, Wysokinski WE, Mauck KF, Bhagra A, Havyer RD, Thompson CA, Hayes SN, McBane RD 2nd. Fibrin d-dimer concentration, deep vein thrombosis symptom duration, and venous thrombus volume. Angiology. 2011;62:253–256. doi: 10.1177/0003319710382416

51. Vucić N, Magdić T, Krnić A, Vcev A, Bozić D. Thrombus size is associated with etiology of deep venous thrombosis--a cross-sectional study. Coll Antropol. 2005;29:643–647.

52. Turitto VT, Hall CL. Mechanical factors affecting hemostasis and thrombosis. Thromb Res. 1998;92:S25–S31. doi: 10.1016/s0049-3848(98)00157-1

53. Cunningham KS, Gotlieb AI. The role of shear stress in the pathogenesis of atherosclerosis. Lab Invest. 2005;85:9–23. doi: 10.1038/labinvest.3700215 [published correction appears in Lab Invest. 2005 Jul;85:942]

54. Davies PF. Hemodynamic shear stress and the endothelium in cardiovascular pathophysiology. Nat Clin Pract Cardiovasc Med. 2009;6:16–26. doi: 10.1038/ncpcardio1397

55. Ghuysen A, Ghaye B, Willems V, Lambermont B, Gerard P, Dondelinger RF, D’Orio V. Computed tomographic pulmonary angiography and prognostic significance in patients with acute pulmonary embolism. Thorax. 2005;60:956–961. doi: 10.1136/thx.2005.040873

56. Irmak I, Sertçelik Ü, Öncel A, Er B, İnam G, Durhan G, Demir A, Çöplü L. Correlation of thrombosed vessel location and clot burden score with severity of disease and risk stratification in patients with acute pulmonary embolism. Anatol J Cardiol. 2020;24:247–253. doi: 10.14744/AnatolJCardiol.2020.55013

57. Beenen LFM, Bossuyt PMM, Stoker J, Middeldorp S. Prognostic value of cardiovascular parameters in computed tomography pulmonary angiography in patients with acute pulmonary embolism. Eur Respir J. 2018;52:1702611. doi: 10.1183/13993003.02611-2017

58. Choochuen P, Kiranantawat N, Nirattisaikul S, Khanungwanitkul K, Chongsuvivatwong V. Can clot size and stenotic degree predict perfusion defects on conventional computed tomographic pulmonary angiography in diagnoses of pulmonary embolism?. Pol J Radiol. 2022;87:e530–e538. doi: 10.5114/pjr.2022.119809

59. Malagò R, Pezzato A, Barbiani C, Alfonsi U, Nicolì L, Caliari G, Pozzi Mucelli R. Coronary artery anatomy and variants. Pediatr Radiol. 2011;41:1505-1515. doi: 10.1007/s00247-011-2218-9

60. Dodge JT Jr, Brown BG, Bolson EL, Dodge HT. Lumen diameter of normal human coronary arteries. Influence of age, sex, anatomic variation, and left ventricular hypertrophy or dilation. Circulation. 1992;86:232–246. doi: 10.1161/01.cir.86.1.232

61. Scarparo P, van Gameren M, Wilschut J, Daemen J, Den Dekker WK, Zijlstra F, Van Mieghem NM, Diletti R. Impact of thrombus burden on long-term clinical outcomes in patients with either anterior or non-anterior ST-segment elevation myocardial infarction. J Thromb Thrombolysis. 2022;54:47–57. doi: 10.1007/s11239-021-02603-3

62. Mirza M, Kummer K, Touchette J, McCarthy R, Rai A, Brouwer P, Gilvarry M. Variability in intracranial vessel diameters and considerations for neurovascular models: a systematic review and meta-analysis. Stroke. 2024;4. doi: 10.1161/SVIN.123.001177

63. Baz RA, Scheau C, Niscoveanu C, Bordei P. Morphometry of the entire internal carotid artery on CT angiography. Medicina (Kaunas). 2021;57:832. doi: 10.3390/medicina57080832

64. Yin Z, Guo J, Li R, Zhou H, Zhang X, Guan S, Tian Y, Jing L, Sun Q, Li G, et al. Common carotid artery diameter and the risk of cardiovascular disease mortality: a prospective cohort study in northeast China. BMC Public Health. 2024;24:251. doi: 10.1186/s12889-024-17749-x

65. Erbel R, Eggebrecht H. Aortic dimensions and the risk of dissection. Heart. 2006;92:137–142. doi: 10.1136/hrt.2004.055111

66. Elefteriades JA, Ziganshin BA, Rizzo JA, Fang H, Tranquilli M, Paruchuri V, Kuzmik G, Gubernikoff G, Dumfarth J, Charilaou P, et al. Indications and imaging for aortic surgery: size and other matters. J Thorac Cardiovasc Surg. 2015;149:S10–S13. doi: 10.1016/j.jtcvs.2014.07.066

67. Hertzberg BS, Kliewer MA, DeLong DM, Lalouche KJ, Paulson EK, Frederick MG, Carroll BA. Sonographic assessment of lower limb vein diameters: implications for the diagnosis and characterization of deep venous thrombosis. AJR Am J Roentgenol. 1997;168:1253–1257. doi: 10.2214/ajr.168.5.9129422

68. Arendt VA, Mabud TS, Jeon GS, An X, Cohn DM 3rd, Fu JX, Hofmann LV. Analysis of patent, unstented lower extremity vein segment diameters in 266 patients with venous disease. J Vasc Surg Venous Lymphat Disord. 2020;8:841-850. doi: 10.1016/j.jvsv.2019.12.078

69. Meissner MH, Manzo RA, Bergelin RO, Strandness DE Jr. Venous diameter and compliance after deep venous thrombosis. Thromb Haemost. 1994;72:372–376.

70. Bosson JL, Riachi M, Pichot O, Michoud E, Carpentier PH, Franco A. Diameters of acute proximal and distal deep venous thrombosis of the lower limbs. Int Angiol. 1998;17:260–267.

71. Tutwiler V, Litvinov RI, Protopopova A, Nagaswami C, Villa C, Woods E, Abdulmalik O, Siegel DL, Russell JE, Muzykantov VR, et al. Pathologically stiff erythrocytes impede contraction of blood clots. J Thromb Haemost. 2021;19:1990–2001. doi: 10.1111/jth.15407

72. Faes C, Ilich A, Sotiaux A, Sparkenbaugh EM, Henderson MW, Buczek L, Beckman JD, Ellsworth P, Noubouossie DF, Bhoopat L, et al. Red blood cells modulate structure and dynamics of venous clot formation in sickle cell disease. Blood. 2019;133:2529–2541. doi: 10.1182/blood.2019000424

73. Samson AL, Alwis I, Maclean JAA, Priyananda P, Hawkett B, Schoenwaelder SM, Jackson SP. Endogenous fibrinolysis facilitates clot retraction in vivo. Blood. 2017;130:2453–2462. doi: 10.1182/blood-2017-06-789032

